# Evidence for continent-wide convergent evolution and stasis throughout 150 years of a biological invasion

**DOI:** 10.1101/2021.04.21.440838

**Authors:** Yihan Wu, Robert I. Colautti

**Affiliations:** Department of Biology, Queen’s University, Kingston, Ontario, Canada

**Author notes:** Botany Department, University of British Columbia, Vancouver, BC, Canada. Corresponding authors Robert I. Colautti and Yihan Wu, **Email:**. **Author Contributions:** Both authors contributed to conceptualization, analysis and writing. In addition, YW wrote the final code and RIC was responsible for funding, supervision and project administration. **Competing Interest Statement:** The authors report no competing interests.

**Keywords:** *Lythrum salicaria*, Virtual Common Garden (VCG), herbarium records, punctuated equilibrium, invasive species, contemporary evolution

## Abstract

The extent to which evolution can rescue a species from extinction, or facilitate range expansion, depends critically on the rate, duration, and geographical extent of the evolutionary response to natural selection. Adaptive evolution can occur quickly, but the duration and geographical extent of contemporary evolution in natural systems remains poorly studied. This is particularly true for species with large geographical ranges and for timescales that lie between ‘long-term’ field experiments and the fossil record. Here, we introduce the Virtual Common Garden (VCG) to investigate phenotypic evolution in natural history collections while controlling for phenotypic plasticity in response to local growing conditions. Reconstructing 150 years of evolution in *Lythrum salicaria* (purple loosestrife) as it invaded North America, we analyze phenology measurements of 3,429 herbarium records, reconstruct growing conditions from more than 12 million local temperature records, and validate predictions across three common gardens spanning 10 degrees of latitude. We find that phenology clines have evolved along parallel climatic gradients, repeatedly throughout the range, during the first century of evolution. Thereafter, the rate of microevolution stalls, recapitulating macroevolutionary stasis observed in the fossil record. Our study demonstrates that preserved specimens are a critical resource for investigating limits to evolution in natural populations. Our results show how natural selection and trade-offs measured in field studies predict adaptive divergence observable in herbarium specimens over 15 decades at a continental scale.

## Introduction

Global biodiversity in the Anthropocene is threatened by the jointly homogenizing effects of population extirpation and biological invasion (1). These outcomes lie along a spectrum of population growth trajectories that depend fundamentally on environmental (mal)adaptation (2). Adaptive evolution in novel and changing environments can rescue populations from extinction (3) and facilitate the spread of invasive species (4). However, genetic constraints can limit the evolutionary response to selection such that the balance of adaptation and constraint within a species determine its ecological niche (5). Species distribution models and management strategies under current and future global change could benefit from a better understanding of the balance between natural selection and genetic constraint, and how this balance affects the rate, duration, and geographical extent of adaptive evolution across a species range.

Selection gradients and rates of phenotypic evolution have been studied in native populations for a variety of taxa (6–9). In contrast, little is known about selection operating in invasive populations, partly due to an apparent researcher bias against measuring selection in non-equilibrial populations (8). More generally, the duration and geographical extent of adaptive evolution are difficult to assess in field experiments, especially when one is interested in variation throughout a species range over decadal to centenary timescales. Common garden studies involving genotypes from contemporary populations can offer insight into local adaptation, but provide only a snapshot of evolutionary change (10–14). ‘Long-term’ ecological studies can reveal temporal dynamics that are not evident in short-term experiments, but are rare due to considerable logistic challenges that also limit the extent of spatial replication (15, 16). Therefore, it remains unclear how the rate and duration of adaptive evolution vary throughout species ranges, and whether evolution can be sustained over the decade to century timescales most relevant to conserve and manage biodiversity for the 21^st^ century.

Invasive species provide opportunities to study evolution in action, but inferences drawn from the genetics of contemporary populations are complicated by demography (i.e. invasion history) and spatio-temporal variation in natural selection (8). Clines in growth and phenology have been identified in many invasive plants, but the rate, duration, and adaptive significance of cline evolution remains uncertain (8, 12). Whether these latitudinal clines are adaptive remains largely untested, but a decade of research has shown that flowering time clines are genetically-based and locally adaptive in invasive populations of *Lythrum salicaria* (purple loosestrife) from eastern North America (17, 18). Local adaptation in these populations likely evolved as a consequence of (i) a trade-off between growth and phenology that is consistent across growing environments, and (ii) directional selection on flowering time with steeper gradients in shorter growing seasons (18–21). This research demonstrates adaptive cline evolution during invasion of *L. salicaria* but covers a relatively small portion of the North American distribution. It is therefore uncertain whether parallel clines evolved elsewhere in North America, and how quickly these adaptive clines evolved during invasion.

Adaptive clines could evolve along two distinct trajectories analogous to long-term evolutionary dynamics observed in the fossil record (Fig. 1). First, a Continuous Evolution Model (CEM) predicts a gradual and relatively constant rate of phenotypic change through time (Fig. 1a,b).

**Figure 1.**
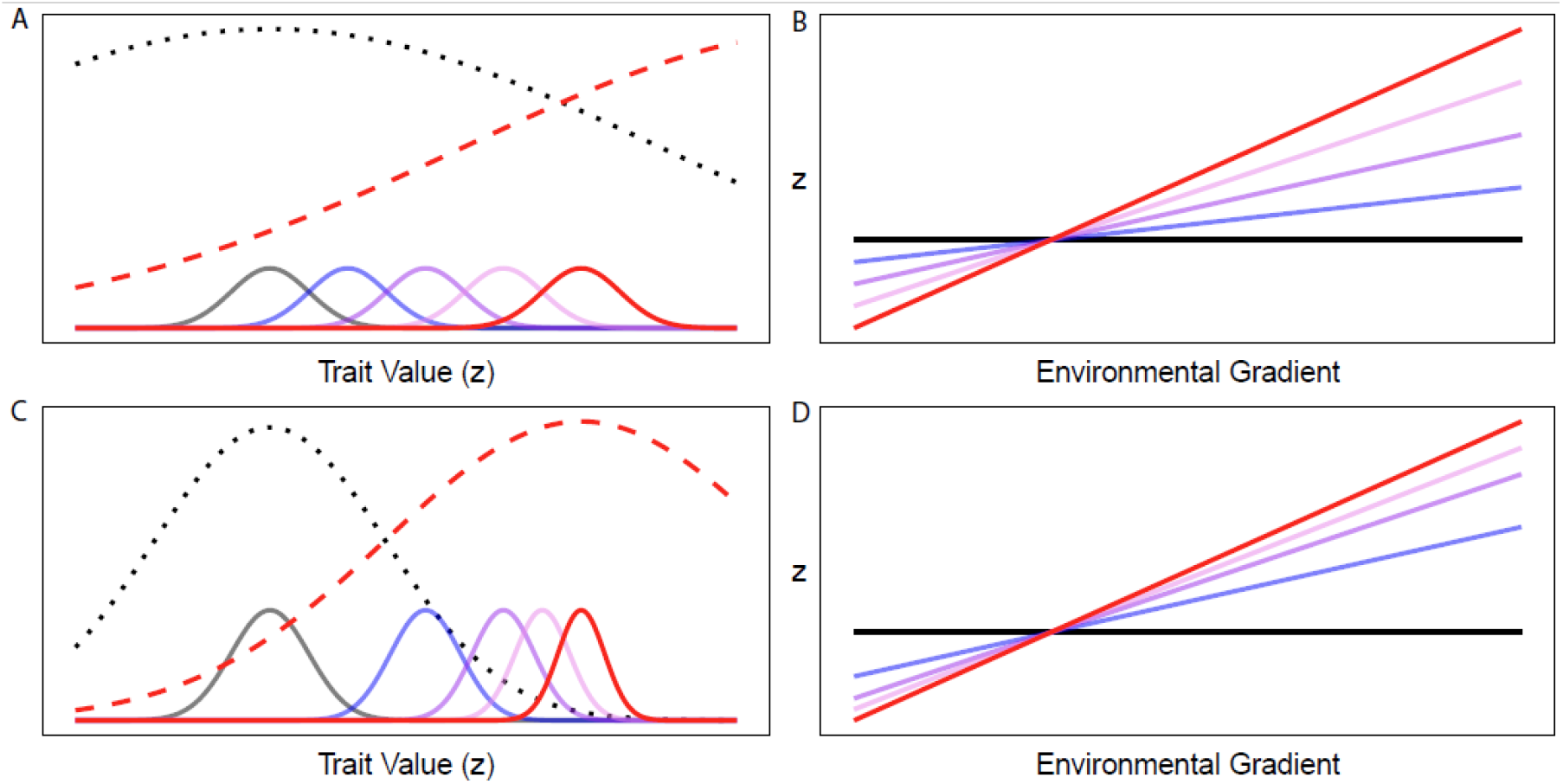
Models of adaptive evolution in a novel environment of a single population (A & C) and multiple populations along an environmental gradient (B & D). In the single population model (A & C), the distribution of phenotypes (*z*) in ancestral populations (solid grey curves) is maintained by stabilizing selection in the ancestral environment (dotted grey curve). Colonization of a novel environment (or a change in local environment) shifts the fitness surface to a new optimum (red dashed curve). A trait under selection evolves towards the new optimum over time (blue through red curves). When adaptive evolution occurs in many populations located along an environmental gradient (B & D), then clines (i.e. slopes) increase in magnitude over time. The Continuous Evolution Model (CEM; A & B) predicts a relatively constant rate of evolution through time. This could occur when (i) stabilizing selection is weak relative to the difference in optima, resulting in approximately linear selection differentials (e.g. dashed red line in A), and (ii) genetic variation introduced by mutation and migration is relatively constant through time. The Punctuated Evolution Model (PEM; C & D) predicts a decelerating rate of evolution. This could occur when stabilizing selection is strong relative to the difference in optima (e.g. dashed red line in C), or when standing genetic variation decreases through time for the trait under selection.

Alternatively, a punctuated equilibrium model (PEM) predicts an early, rapid burst of evolution followed by a static equilibrium (22). These scenarios represent opposite ends of a spectrum defined by (i) the strength of natural selection and (ii) the amount of genetic variation for the trait under selection (23). In the CEM, evolution is relatively slow and approximately continuous because the change in fitness across environments is approximately linear and genetic variation is not limiting. In the PEM, a punctuated response to evolution occurs when there is strong directional selection initially, but populations quickly reach their new optima and/or standing genetic variation is depleted during the response to selection. Distinguishing which models apply to natural populations is important for understanding the rate of contemporary evolution in response to novel and changing environments. Yet, testing these alternatives across a species range requires quantifying rates of evolutionary change at multiple locations.

Temporal sampling of genetic changes in natural populations provides some of the strongest evidence for adaptive evolution in nature (16, 24), but these experiments are difficult to replicate over large spatial and temporal scales (25–27). In contrast, millions of preserved specimens maintained in global natural history collections offer detailed records of phenotypes sampled over decades to centuries past, holding potential for reconstructing evolutionary trajectories (28–30). Indeed, herbarium records can be used to reliably reconstruct phenology variation over space and time (31), but evolutionary shifts in phenotypic traits observed under natural field conditions are difficult to distinguish from developmental plasticity associated with different environments (32). Thus, it would be valuable to develop a mechanistic model, informed by field experiments, to disentangle phenotypic plasticity from genetic effects on phenotypes observed in natural history collections. Such a model could enable tests of adaptive evolution and genetic constraint across broad temporal and spatial scales unachievable by other methods.

The virtual common garden (VCG) is a computational approach that incorporates biophysical models informed by field experiments to reconstruct phenotypic plasticity and genetic differentiation over fifteen decades at a continental scale. As such, the VCG is a type of observational experiment for eco-evolutionary inference. Using this approach, we reconstruct 150 years of phenotypic evolution in the self-incompatible wetland perennial *Lythrum salicaria* (purple loosestrife) as it spread across North America. Our specific goals are: (i) to parameterize and then validate the VCG as a method to distinguish genetic variation from developmental plasticity in *L. salicaria*; (ii) to test for convergent evolution via local adaptation across North America; (iii) to test the CEM and PEM models shown in Figure 1 by measuring changes in the rate of cline evolution through time.

## Results

### Herbarium Specimens

We calculated the spatially-weighted interpolation of Growing Degree Days up to the date of specimen collection (*GDD*_*C*_) and the total growing Season Length (*SL*) for each of the 3,429 digitized herbarium specimens. These were based on >12 million temperature records, used to reconstruct 62,208 annual temperature accumulation curves from 6,303 weather stations (Fig. 2).

**Figure 2.**
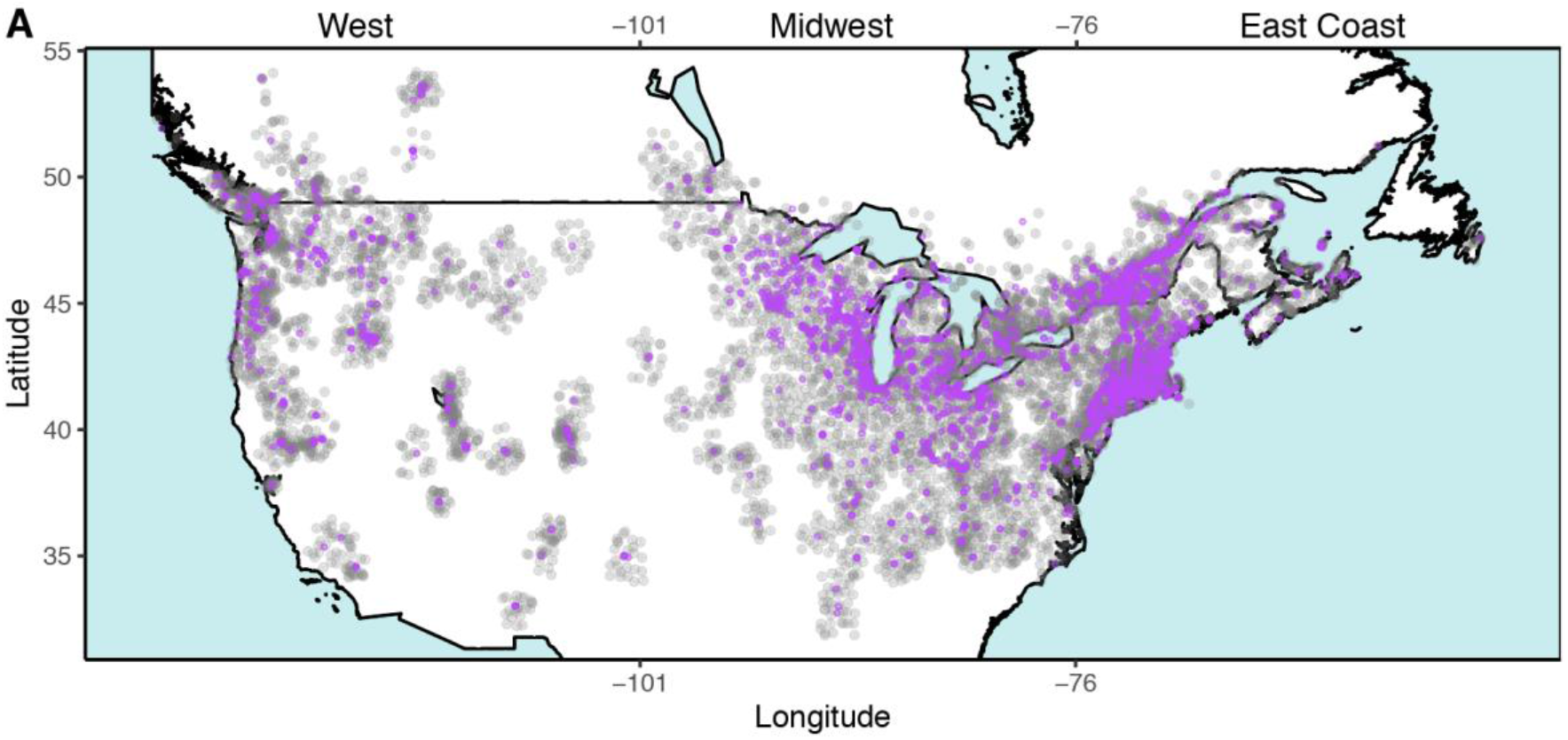
Map of the United States and Canada showing locations of 3,429 *Lythrum salicaria* specimens (purple dots) collected between 1866 and 2016, and locations of 6,303 weather stations (light grey dots) used to interpolate local climate data. Specimens are classified into three geographical regions with different invasion histories: West (< -101° longitude), Midwest (−76° to -101°), and East Coast (> -76° longitude).

Measurements from all digitized herbarium specimens that met inclusion criteria (Fig. 2) are available online in the Dryad Database (https://doi.org/10.5061/dryad.qfttdz0f5) with images available from sources cited therein. The earliest specimen was collected on July 29th, 1866 near Boston, Massachusetts, with the number of samples increasing thereafter to a peak in the 1970s and declining in recent decades (Fig. S2). The most recent specimen was collected on August 31, 2016 from Lake Lowell in Idaho. Specimens cover a large part of North America spanning from Atlantic to Pacific coasts, and almost 20 degrees of latitude, from as far south as Carthage, Mississippi, USA (32.74°N) to Prince George in BC, Canada in the north (54°N). A majority of specimens were collected in the East Coast and Midwest regions (Fig. 2), each with over 1,000 specimens, while the West region contains approximately 400 specimens (Table S1). The most densely sampled regions are the Great Lakes region (particularly in Illinois and Wisconsin), along the St. Lawrence River, and along the northeastern coastal region of North America (Fig. 2).

The average Julian day of collection was 217 d (*sd* = 25.18), or August 5 in a non-leap year, and ranged from 121 d (May 1, 1938) to 365 d (December 31, 1906). This represents a broad range of days measured from the start of the growing season at each location, with a mean of 98 d (min = 8, max 263) (Fig. S4). Growing season length (SL) estimated from weather stations varied across regions, with the average growing season longer in the West Coast region and decreasing eastward; SL was also strongly correlated to latitude (*r* = -0.6). Specimens captured a broad distribution of phenological stages from a few buds to fully mature fruits. There was no clear evidence for bias with respect to phenology sampling, as all but the earliest phenologies were evenly distributed (Fig. S3). Nor did we find strong clustering of phenologies with respect to geography, even though some structure is expected due to the predicted relationship between season length and latitude (Fig. S5). However, we did observe variation in sampling intensity across the range (Fig. 1), which we address using spatially-explicit resampling methods as described later.

### VCG Validation

Details of the VCG analysis are provided in the Materials and Methods, Supplementary Methods and online repository (DOI) (https://doi.org/10.5061/dryad.qfttdz0f5). The VCG analysis involves several key steps, which are outlined in the Materials and Methods section.

To validate the VCG we compared (i) latitudinal clines in development time (*ψ*) of herbarium specimens with (ii) days to first flower in three real-world common garden experiments in eastern North America (data from (18)), each standardized to a mean of zero and unit variance. This comparison involves heterogenous phenology measurements (i.e. *ψ* and days to first flower) and growing conditions (i.e. a virtual common garden and three ‘real world’ garden sites spanning 10° of latitude), yet the estimated latitudinal clines were remarkably consistent (Fig. 3). The correlation coefficients (i.e. clines) observed in the VCG (*r* = -0.70, bootstrapped 95% CI: - 0.17 to -0.92) was not significantly different from those measured in real-world field transplant sites at northern (*r* = -0.84, CI: -0.70 to -0.95), mid-latitude (*r* = -0.77, CI: -0.60 to -0.90) and southern gardens (*r* = -0.87, CI: -0.75 to -0.95).

**Figure 3.**
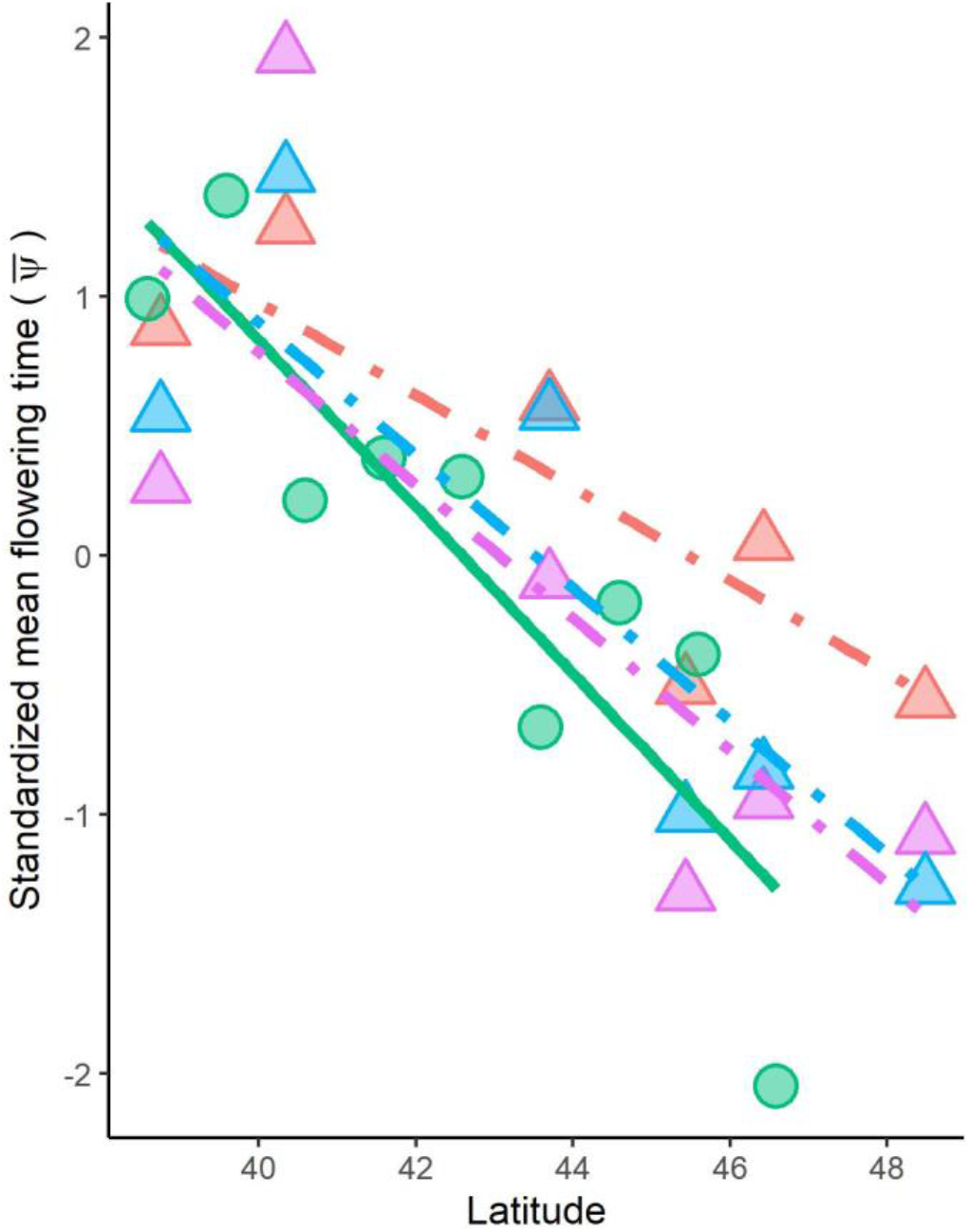
Validation of clines observed in the Virtual Common Garden (VCG). Standardized estimates (z-score) of mean phenological development time 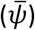 were inferred from contemporary herbarium specimens from eastern North America, binned by one degree of latitude (green circles). This cline is similar to average z-scores of flowering times estimated for six populations grown at each of three reciprocal transplant sites spanning 10° of latitude: Timmins, Ontario (blue triangles), Koffler Scientific Reserve near Newmarket, Ontario (purple triangles), and Blandy Experimental Farm near Boyce, Virginia (red triangles).

### VCG Cline Evolution

Clines in phenology were observed in all three geographic regions for specimens collected after 1980, and generally weaken back through time (Fig. 4, Table S3). East Coast populations were already well-established long before the first specimen collection in 1866 (33), and show a relatively stable cline until the present day. In contrast, contemporary populations in the Midwest have a similar cline that weakens back toward the time of establishment (∼1900), while West Coast populations did not establish until the 1920s and have not reached the same magnitude of cline as the two older regions (Fig. 4).

**Figure 4.**
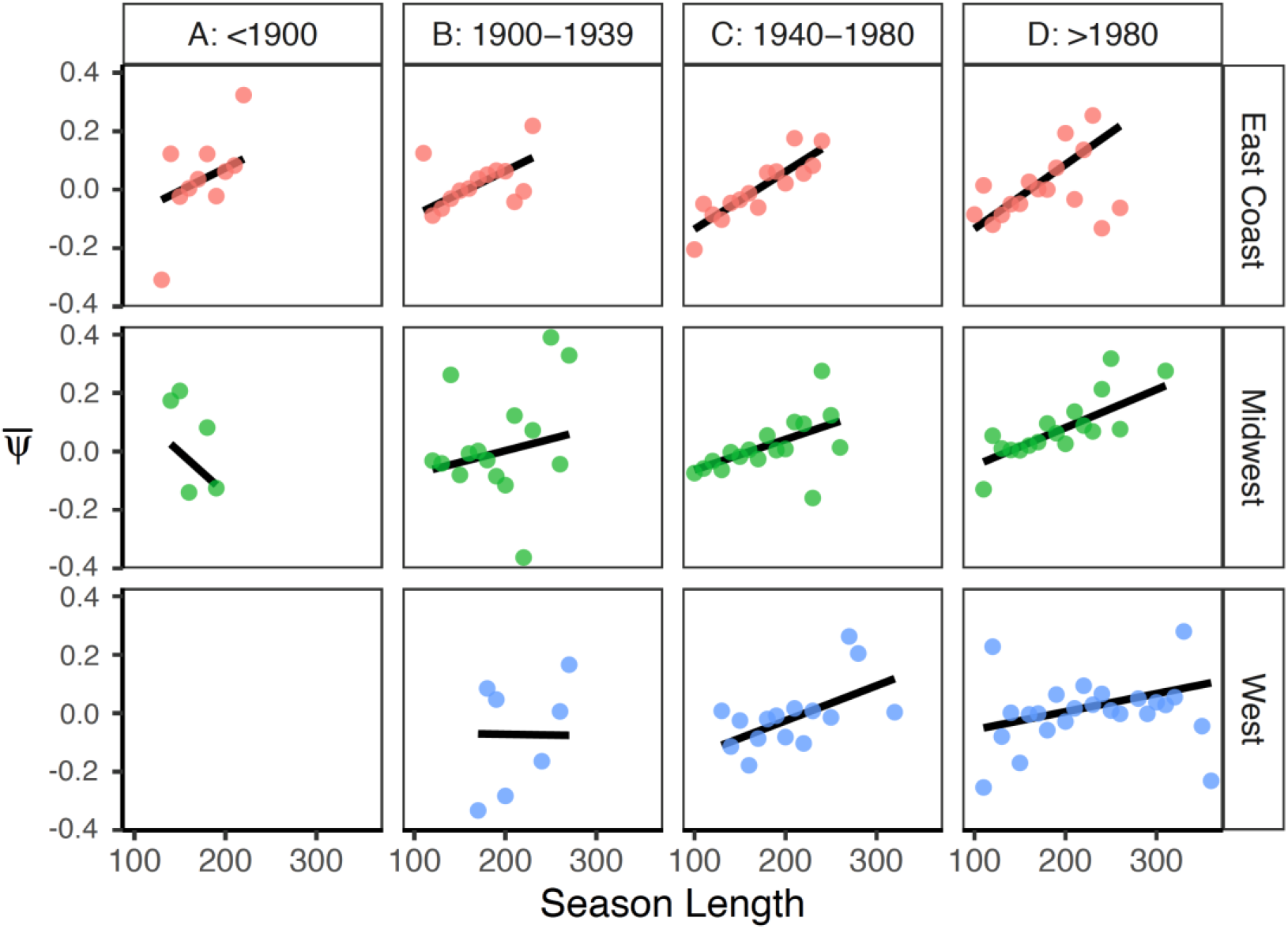
Spatial (rows) and temporal variation (columns) in phenological clines of *Lythrum salicaria* as it spread across North America. Clines in the East Coast (red dots, top row), Midwest (green dots, middle row), and West regions (blue dots, bottom row) are divided by time of collection, binned into four eras (columns). Each bivariate plot shows the relationship between the mean phenological development time of herbarium specimens 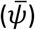 and the mean estimated season length, averaged across specimens binned every ten days of season length.

As a formal statistical test for a change in evolutionary rates across North America, we first modelled phenological development as a sigmoidal function of accumulated growing degree days (*GDD*_*c*_). Analogous to the traditional common garden experiment, this model attempts to minimize phenotypic variance associated with developmental plasticity. To test for local adaptation, we add a parameter for season length (*SL*) as the predicted genetic correlation between season length and phenology. Parameter estimates for *GDD*_*c*_ and *SL* were significantly different from zero, and there was a significant *GDD*_*c*_ × *SL* interaction (Table S4). Together, these parameters represent a shift in phenology curves (Fig. 5a) with southern genotypes (red curves) predicted to take longer (x-axis) to reach a particular phenological stage (y-axis). If days to first flower is mapped to the x- and y-axes in Fig. 5a, the model recovers the common garden clines shown in Figure 3 in which genotypes from southern locales with longer growing seasons flower later than genotypes from shorter growing seasons.

**Figure 5.**
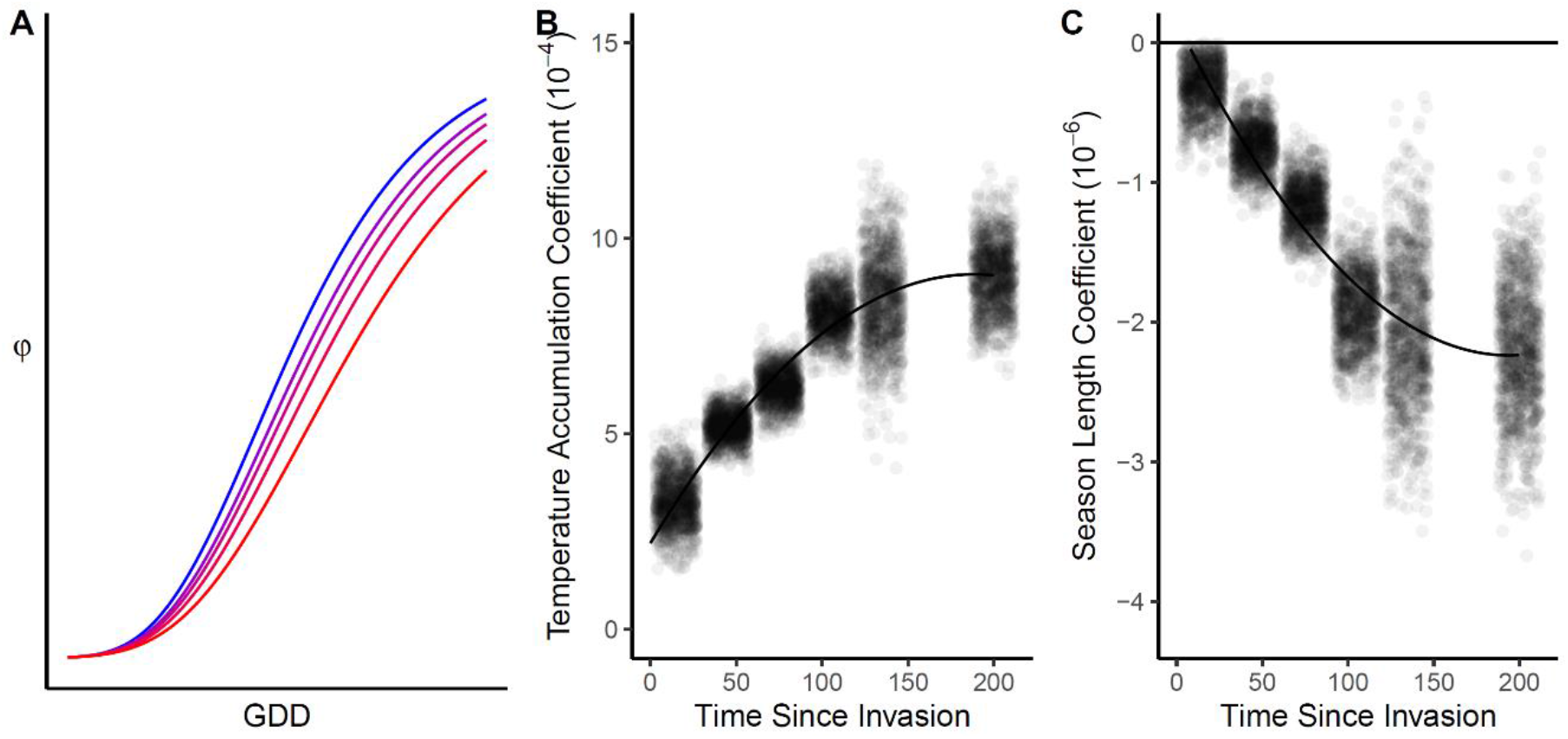
A nonlinear least-squares (NLS) model testing temporal changes in phenology clines. (A) A model of observed phenological stage (*φ*) as a function of local growing conditions (growing-degree days at the time of collection; *GDD*_*C*_) for five different season lengths (*SL*) parameters ranging from short/northern (blue) to long/southern (red). (B) Bootstrap estimates of *β*_*b,t*_, the model coefficient for *GDD*_*C*_. (C) Bootstrap estimates of *β*_*d,t*_, the model coefficient for season length (*SL*). Bootstrap estimates include 1,000 spatially explicit resampling iterations (with replacement) with specimens binned into one of six classes of population age (i.e. time since establishment). Lines in B and C show quadratic linear regression of bootstrap estimates.

In addition to testing for phenology clines, we used the same model to estimate evolutionary rates. Using a spatially explicit bootstrap resampling model to estimate *SL* and *GDD*c parameters over time, we found that the inferred rate of evolution was significantly different from zero (non-zero slope), and decelerating (significant quadratic term), rather than remaining constant (linear slope) during invasion (Fig 5c). Specifically, the coefficient estimates for linear and quadratic models are reported in Table S5 for the effect of temperature accumulation on phenology (*β*_*b,t*_), and Table S6 shows the coefficients for the effect of season length on phenology (*β*_*d,t*_). Based on Likelihood Ratio Tests (LRT), the quadratic terms were significant for both *β*_*b,t*_ (*χ*^*2*^ = 5448, p < 0.001) and *β*_*d,t*_ (*χ*^*2*^ =3045, p < 0.001), with a ∼15% increase in *R*^*2*^ values (*β*_*b,t*_: 0.763 to 0.876; *β*_*d*_: 0.713 to 0.810). This is shown in Figure 5 wherein the coefficients for *β*_*b,t*_ (Fig. 5b) and *β*_*d*_ (Fig. 5c) stabilize around 100 years post-establishment.

## Discussion

Geographical clines in morphological, physiological, and life-history traits are common to many taxa, but it has been difficult to measure the rate, duration, and geographical extent of cline evolution occurring in natural populations. In this study, we introduced the Virtual Common Garden (VCG) analysis as a multi-step computational approach to test evolutionary hypotheses using natural history collections. Below, we explore the relevance of our findings to the spread of invasive species and more generally to predict adaptive evolution in novel and changing environments.

### Convergent Evolution During Invasion

Feedbacks between colonization and trait evolution can facilitate rapid range expansion (4, 34, 35), but clines *per se* do not provide clear evidence for adaptive evolution. In particular, simulations show how non-adaptive and maladaptive clines may evolve frequently when repeated founder events occur during spread, producing statistically significant but random slopes throughout the range (8, 36). In the case of *L. salicaria*, reciprocal transplant experiments spanning > 1,000 km and five growing seasons confirm that populations are locally adapted in eastern North America, with adaptive phenologies boosting reproductive fitness by one to two orders of magnitude (18). Heretofore, it had been unknown whether natural selection was likely a causal agent of cline formation or merely acted in a supporting role, maintaining clines that evolved primarily through stochastic processes. Our VCG results do not support the prediction of random clines produced by founder effects and drift. Rather, similar clines evolved repeatedly throughout the range and strengthened over time (Fig. 4), implicating adaptive evolution as an important driver of phenology evolution across North America. These replicate, adaptive clines offer a case-study in intraspecific convergent evolution of a quantitative trait.

Classic examples of convergent evolution typically focus on distantly related species with similar but derived adaptations. In contrast, intraspecific convergent evolution of a phenotypic trait is not well-defined (37–39). In our analysis, phenologies of geographically isolated populations converge when they experience similar season lengths, while adjacent populations diverge when season lengths differ (Figs. 4 and 5). Regardless of the relative contribution of ancestral and derived alleles, adaptive evolution across a heterogenous adaptive landscape should leave a signature of phenotypic convergence, which we observe in the herbarium record as a reduction in spatial entropy of *ψ* over time. Intraspecific convergent evolution via local adaptation has implications for predicting the fate of introduced and native species in the Anthropocene.

### Adaptation, Constraint and Stasis

Rapid evolution can facilitate persistence and spread in novel and changing environments, but trade-offs may constrain the rate and duration of adaptive evolution. We therefore hypothesized that adaptive clines could evolve along two distinct trajectories analogous to long-term evolutionary dynamics observed in the fossil record (Fig. 1). Our analysis supports the Punctuated Equilibrium Model (PEM) over the Continuous Evolution Model (CEM), with a rapid but relatively constant rate of cline evolution for ∼100 years followed by a contemporary period of relative stasis (Fig. 5). Although several different scenarios of natural selection and genetic variation could produce a decline in evolutionary rates consistent with the PEM (Figs. 1 and 5), previous studies of *L. salicaria* suggest trade-offs as a primary factor.

Multiple common garden studies of *L. salicaria* from eastern North America reveal ample genetic variation within and among populations, measured for both molecular markers and quantitative traits (17, 20, 40). Populations also experience strong directional selection for flowering time, growth, and related quantitative traits (18, 19). Yet, despite significant standing genetic variation and strong directional selection, local adaptation is maintained by antagonistic selection on correlated traits involved in growth and phenology (18, 21). A punctuated evolution of clines followed by the stabilization of evolutionary rates consistent with PEM add evidence that this adaptation-constraint phenomenon may be relevant to adaptation throughout a species range. Specifically, adaptive evolution should be fastest in the early stages of colonization when newly established populations are far from their phenotypic optima and thus experience stronger directional selection (Fig. 1). Evolutionary rates should decline thereafter as antagonistic selection on plant size maintains the local optima, resulting in a static equilibrium that is not observed for the first several decades of invasion. It is an open question as to whether this phenomenon is common among biological invasions and more generally whether rates of evolution measured in contemporary populations will stall over longer timescales.

An inherent assumption in the application of a VCG analysis is that plasticity of a trait of interest (e.g., phenological development time) can be predicted by local growing conditions (e.g., *GDD*_*c*_), with limited genotype-by-environment interactions (G x E). Another limitation is that the model cannot parse unmeasured environmental characteristics (*U*_*N*_) (e.g., photoperiod), which could confound our interpretation if the *U*_*N*_ affect growth and correlate with *SL*, but do not similarly correlate with *GDD*_*c*_. Multiple common garden experiments with *L. salicaria* confirm that clines in growth and phenology are similar across rearing environments with limited influence of G × E or photoperiod effects (17–19, 41). Moreover, confounding effects of G x E or *U*_*N*_ would have been evident as non-parallel clines in the common garden data shown in Figure 3. Instead, we observe a constancy of clines across rearing environments, both real and virtual.

A pattern of rapid evolution and stasis may be common during invasion, given that the climatic niche is often conserved between the native and introduced range (42). Similar to *L. salicaria*, many other invasive species also (i) exhibit genetically-based clines, (ii) experience directional selection, and (iii) segregate genetic variation for quantitative traits at levels comparable to genotypes from their native range (8, 12). It is unclear whether native species would show similar dynamics during natural colonization (e.g. post-glaciation), as the genetics of native populations can be quite different (2). Nevertheless, our study demonstrates how genetic constraints can help predict evolutionary outcomes that may take decades to manifest. A key goal for future VCG studies is to identify genetic clines and differentiate those that evolved through stochastic processes (e.g. random clines) vs. natural selection (e.g. convergence). Convergent cline evolution and stasis could provide a foundation for investigating constraints on evolution in novel environments that might not be evident for the first few decades of evolution.

### Natural History Collections and Evolution

Studies of evolutionary change in natural populations often rely on demographic inferences from molecular markers, but inferences from allele frequency data are known to be problematic for genetic loci of quantitative traits like phenology, and for populations with complex demographic histories that have not reached an equilibrium between migration/mutation and drift/selection (8, 43, 44). For example, a shift from strong directional to weak stabilizing selection on a set of multi-locus traits following the PEM scenario in Figure 1 could leave a genomic signature similar to a demographic bottleneck followed by population expansion. These limitations help to explain why Q_ST_-F_ST_ analyses of *L. salicaria* populations do not reveal selection on growth and phenology (40), contrary to clear evidence in field studies that these traits are under strong selection (18, 19, 21). In contrast, our VCG analysis identified convergent clines that are consistent with locally adaptive clines confirmed in field experiments (Fig. 3). Additionally, the VCG revealed evidence for repeated evolution of clines across decades and thousands of kilometers (Fig. 4), with variable rates of change (Fig. 5). Although inferences from neutral genetic markers in contemporary populations may be limited, the application of molecular markers in combination with herbarium records could help to overcome some of these inherent limitations.

Informed by prior empirical studies and trade-off models, convergent cline evolution and stasis inferred from herbarium records of *L. salicaria* demonstrate how evolution of ecologically important traits in natural systems can be predictable on scales most relevant to modern human civilization: continental to global spatial scales and decadal to centenary timescales. A VCG analysis of herbarium specimens is just one example of how natural history collections can be mobilized to complement limitations of scale and simplism necessary for tractable experimentation. Preserved specimens offer a rare opportunity to rewind the clock and reconstruct phenotypic change, but they are vulnerable to loss without significant financial and academic investment. Global efforts to manage biodiversity in the Anthropocene could benefit from a more predictive science of adaptive evolution and constraint in natural systems, made possible only with the support of natural history collections at a global scale.

## Materials and Methods

### Virtual Common Garden

Common garden studies are commonly used to identify and quantify genetic differences in quantitative traits. Genetic variation is inferred from phenotypic differences among populations, genetic families, or genetic clones when all individuals are reared under similar growing conditions. The Virtual Common Garden (VCG) applies the same philosophy but uses a computational approach that includes the following steps:

1. Convert inflorescence measurements of each specimen to a size-independent and scale-free index of phenological stage (*φ*).
2. Generate histograms of *φ* to look for sampling bias.
3. Map the geographic location and phenological stage of each specimen to look for spatial bias.
4. For each specimen, download weather data for up to 20 nearby weather stations.
5. For each weather station, calculate a vector of 365 (or 366) average daily temperatures for the same year in which a nearby specimen was collected.
6. Interpolate a vector of average daily temperature for each specimen by inverse distance weighting.
7. Estimate the start and end of the growing season for each specimen to define the local season length (SL).
8. Calculate the local growing-degree days from the start of the growing season to the day of collection (*GDD*_*c*_) for each specimen.
9. Fit a statistical model of observed phenological stage (*φ*) and collection date to estimate a phenological development time (*ψ*) for each specimen.
10. Use spatial Kriging to interpolate population time since establishment for each specimen (Age). Using these data, the VCG does the following:
11. Validate *ψ* by comparison with flowering times observed in common garden experiments.
12. Calculate correlation between *ψ* and season length, and test whether these change over space and time.
13. Use another statistical model that will (i) predict the joint effects of *GDD*_*c*_ and local adaptation to season length, and (ii) test whether these relationships (i.e. model coefficients) change over time.
14. Resample, with replacement, individual specimens at 0.1 ° latitude/longitude resolution to generate bootstrap means and confidence intervals for the model parameters.

### Phenology of Herbarium Specimens

Between August 2016 and December 2017, we analyzed 3,429 digitized herbarium specimens obtained from five sources: (i) the Global Biodiversity Information Facility (GBIF, http://www.gbif.org/), (ii) the Regional Networks of North America Herbaria accessed through the Arizona-Mexico Chapter (http://swbiodiversity.org), (iii) the New York Botanical Garden (NYBG, http://www.nybg.org/), and (iv) the Database of Vascular Plants of Canada (VASCAN) (*20*)(45) and v) correspondence with 19 university herbaria to obtain more images (see acknowledgements). We included only specimens with both a full collection date (year, month, day) and location information that could be georeferenced (see Supplementary Methods).

Inflorescence development in *L. salicaria* occurs acropetally from the base of the stem to the tip of the apical meristem, resulting in three distinct regions of the inflorescence: fruits (basal), flowers, and buds (distal). To capture variation in floral phenology, we measured the length of each segment on each specimen using the segmented tool in ImageJ (46). We did this for the primary meristem, unless it was clearly damaged in which case the longest inflorescence was measured. Unpollinated flowers occur early in the phenology when compatible mates or pollinators are unavailable. As a result, they senesce and fall from the stem, leaving distinctive callouses at the point of attachment that represent the same phenological stage as fully-formed fruits. Using these measurements, we calculated a scale-free phenological stage (*φ*) for each herbarium specimen *i*:

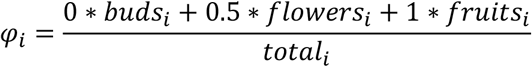

Values of *φ* range from 0 (early) to 1 (late) as a measure of phenological stage at the time of collection. Thus, on any given collection date, specimens with a phenology stage closer to 0 represent phenotypes early in their phenology whereas those with a stage closer to 1 represent phenotypes sampled later in phenological development.

Herbarium specimens are not often random samples of natural populations. Instead, sampling locations are chosen deliberately or opportunistically, introducing potential sampling biases. For example, first flowering dates observed in herbarium specimens collected from New England (USA) were biased toward a delay of ∼3d relative to field observations; however dates were highly correlated with field observations despite this bias (31). Spatial and temporal clustering of samples is also likely when multiple specimens are often collected in sampling excursions conducted by a single individual or group. We therefore developed an analysis that would be robust to phenological stage, sampling date, absolute flowering date, and spatial clustering, as described below.

### Season Length and Growth Conditions

Season length (*SL*) is a key predictor of genetic differentiation for flowering phenology in *L. salicaria*, whereas growing-degree days at the time of collection (*GDD*_*C*_) determines the rate of growth and development and thus plasticity in phenology (18, 47–49). Both *SL* and *GDD*_*C*_ were interpolated from computation of daily weather data available from the Global Historical Climatology Network-Daily database (50). The algorithms and code for the computational interpolation are available on the Dryad Database (https://doi.org/10.5061/dryad.qfttdz0f5). Briefly, we collected temperature data for each herbarium specimen from up to 20 of the closest weather stations located within 0.5 ° latitude and longitude (Fig. 2). For each weather station, we identified the first interval of at least 10 consecutive days above a threshold growing temperature (8 °C) from Jan 1 in the year of collection. The first day of this interval was set as the start of the growing season, and the end of the growing season was determined as the first day thereafter in which the temperature fell below 8° C. Temperature accumulation above 8° C is a key factor affecting growth and development of *L. salicaria*, and commonly used in growth models for the species (17, 47, 49, 51). A length of 10 days at the start of the season was chosen to avoid shorter intervals of abnormally warm winter or spring temperatures where growth would be unlikely. We then used the season start date to calculate the cumulative growing degree days until the day of collection (*GDD*_*C*_). In summary, *GDD*_*C*_ characterizes the local growing environment in order to correct for variation in growing conditions and sampling date, whereas variation in season length (*SL*) is predicted to drive differentiation via local adaptation to climate.

### Phenological Development Time

Field surveys and common garden experiments with *L. salicaria* have demonstrated both plastic and genetic variation for flowering phenology – warmer temperatures accelerate phenological development, whereas natural selection has caused the evolution of earlier flowering in short growing seasons (18, 47). By comparison, population-by-environment interactions are relatively weak, even across field sites spanning a latitudinal gradient of 1,000 km (18). Leveraging these observations, the VCG uses temperature accumulation data to model phenology as the additive effects of genotype and environment (plasticity). Since we cannot retroactively grow herbarium specimens in a common garden, we instead correct the observed phenological stage of each specimen (*φ*_*i*_) for variation in the local growing environment (*E*_*i*_) to estimate a relative measure of phenological development time due to genetic factors (*ψ*_*i*_):

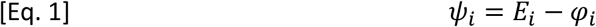

As described later, *E*_*i*_ is the predicted contribution of environment to phenotype in a nonlinear least-squares model of phenology as a function of *GDD*_*C*_. The equation for phenological development time (*ψ*) is therefore the residual phenological stage after accounting for *GDD*_*C*_, multiplied by negative one. We use the negative multiple so that values for phenology in herbarium specimens lay in the same direction of more common phenological measurements (e.g. days to first flower, time to maturity).

In summary, we use two indices to describe the phenology of each herbarium specimen: The phenological stage (*φ*) is calculated directly from inflorescence measurements. The phenological development time (*ψ*) is an estimate of relative phenology after controlling for local growing environment. Phenological stage (*φ*) ranges from 0 (early stage) to 1 (late stage) and is determined by genetic factors, local growing conditions, and collection date. In contrast, the phenological development time (*ψ*) ranges from -1 (fast phenology) to 1 (slow phenology) and predicts genetic effects on phenology. As such, *ψ* is a standardized phenology metric that predicts which phenotypes would be observed when grown under similar rearing conditions – a virtual common garden.

### VCG Validation

To validate the VCG, we compare the predicted phenological development time (*ψ*) inferred from herbarium specimens with observed flowering time clines reported in three separate common garden field experiments. These experiments are part of a previously published reciprocal transplant study involving six populations sampled along a gradient of 10° latitude in eastern North America and grown at three sites: Timmins, Ontario (TIM), the Koffler Scientific Reserve (KSR) near Newmarket, Ontario, Canada, and the Blandy Experimental Farm (BEF) near Boyce, Virginia (18).

To facilitate comparison of virtual and real common garden metrics, we standardized population means within each dataset to z-scores (Mean = 0, SD = 1) and included only herbarium specimens collected after 1960 from the same geographic range in northeastern North America. Note that we standardize population means, rather than individual specimens, because population means are the unit of comparison between herbarium samples and common garden plants for validation purposes. We chose 1960 as a cutoff to maximize sample size while reducing the influence of early collections that could have occurred long before populations became locally adapted. However, we also examined temporal changes in phenology, as described in the next section.

Herbarium specimens were binned into geographic populations at intervals of 1° latitude for comparison with geographic populations sampled in the reciprocal transplant experiment. For each of the four ‘gardens’ (i.e. VCG, TIM, KSR, BEF) we calculated bootstrap estimates (1000 iterations each, with replacement) of latitudinal clines and 95% confidence intervals. In each iteration, we calculated the correlation between the population mean z-score and latitude, resampling either the phenological development time (*ψ*) of herbarium specimens (*N* = 449) binned into nine geographic ‘populations’ (VCG) or average flowering times of seed families sampled from six populations (*N* = 82).

### Population Age and Cline Evolution

After validating the VCG analysis for eastern North America, we tested for parallel latitudinal clines predicted by a selection-constraint model of local adaptation to season length (18, 20, 21). This model predicts an evolutionary shift from slow to fast phenology under shorter season lengths. We also tested the CEM and PEM predictions (Fig. 1) by looking at temporal changes in the rate of cline evolution. In addition to examining clines in season length in a combined statistical model, we separately analyzed three regions with different invasion times that are also separated by natural gaps in the distribution (Fig. 2): East Coast (< 76 °W), Midwest (76 °W-101 °W), and West Coast (> 101 °W).

To examine changes in the rate of evolution over time, we first had to estimate population age, which we did using Kriging interpolation in R (52) (Fig. S1). In addition to collection dates of available herbarium records, we also included dates of other field observations available from the Global Biodiversity Information Facility (GBIF), Biodiversity Information Serving Our Nation (BISON), and Vascular Plants of Canada (VASCAN) (45, 53, 54). The resulting 11,735 records were pooled into grid units of one degree latitude and longitude, and within each grid unit, the year of the earliest observation was used to estimate the date of colonization. Our estimate of population age for each specimen was calculated by subtracting the collection year of each specimen from the locally interpolated (i.e. Kriged) year of invasion.

### Statistical Methods

We modeled the observed phenological stage of each herbarium specimen (*φ*_*i*_) using a logistic model (Eq. 6 in (55)), and a more detailed nonlinear least squares (NLS) regression equation based on the Unified Gompertz curve (Eq. 8 in(55)). We report the Gompertz model here but the logistic model produced similar results (see Supplementary Methods). The analysis is based on a sigmoidal curve of the form:

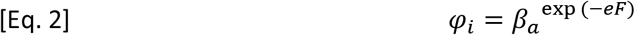

where *β*_*a*_ is an estimate of the y-intercept and *F* is a statistical model that differs for each of three analyses, as outlined below.

First, we calculated a standardized metric of phenology development time for each herbarium specimen (*ψ*_*i*_). This metric enables comparison with genetic differences observed in common garden experiments, and it can be used to visualize genetic changes over space and time. To calculate *ψ*_*i*_, we modeled the observed phenological stage (*φ*_*i*_) as a function of growing environment (*GDD*_*C*_) using a variant of Equation 2 in which

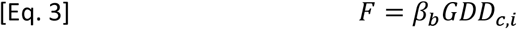

Predictions from this model are the predicted contribution of growing environment to phenology (*E*_*i*_) shown in the VCG analysis described in Equation 1. Subtracting the observed phenological stage from the stage expected for a given *GDD*_*c*_ yields an estimate of the phenological development time (*ψ*_*i*_). This is mathematically equivalent to taking the residuals of Equations 1 & 3 and back-transforming to the scale of *φ*_*i*_.

In a second statistical model, we simultaneously estimated the relative effects of growing environment (*GDD*_*C*_), season length (*SL*) and population age (*Age*). We also examined how the effects of *GDD*_*C*_ and *SL* change with population age of each herbarium specimen:

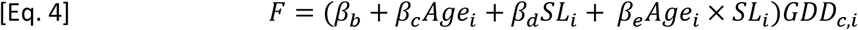

We refer to *β*_*b*_ as the temperature accumulation coefficient, *β*_*c*_ as the population age coefficient and *β*_*d*_ as the season length coefficient. The coefficient *β*_*e*_ enables a statistical test of the hypothesis that clines strengthen over time, while *β*_*c*_ tests the hypothesis that cues evolve to better predict growing conditions.

To avoid problems with spatial heterogeneity in the collections (e.g. pseudoreplication due to geographical sampling bias), we calculated bootstrap means and 95% Confidence Intervals by randomly resampling, with replacement, a single specimen within each cell of a 0.1° latitude by 0.1° longitude grid (N = 1,000 iterations).

Our third statistical model was applied to specimens binned into six age classes: < 30 years, 31-60, 61-90, 91-120, 121-150, and > 151. Within each age class, we used the 0.1° resampling bootstrap model as above. Without population age as a predictor, the equation for each bin simplifies to:

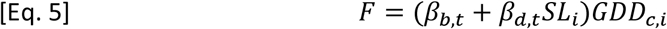

To test the CEM and PEM predictions, we fit linear and quadratic models to the bootstrapped estimates of *β*_*b,t*_ and *β*_*d,t*_ in Equation 5, using the median age of each time bin (*t*) as the predictor variable and the model coefficients as a response variables.

## Acknowledgments

We thank SCH Barrett, LH Rieseberg, CA Rushworth, KG Turner, an anonymous handling editor and three anonymous reviewers for manuscript feedback, and the following people for generously digitizing herbarium records for our study: M Pace & B Thiers (NY Botanical Garden), D Brandon (U Memphis), CA McCormick (UNC Chapel Hill), J Shepard & T Barry (UC Davis), L Struwe (Chrysler Herbarium), R Kleinman (Zimmerman Herbarium SNM), M Nazaire (Rancho Santa Ana Botanic Garden), A Sanders (UC Riverside), P Wolf (Utah State U), D Stover (Kent State U), D Fabijan (U Alberta), S Perkins (San Juan College), K Damboise (Herbier Louis-Marie), N Cowden (Ramsey-Freer Herbarium), S Fuentes-Soriano (New Mexico State U), W Erica (Royal Museum of British Columbia), D Giblin (U Washington), M Hooker (Marion Ownbey Herbarium), A Liston (Oregon State U), A Lopez Villalobos (Fowler Herbarium). This research was supported by NSERC Discovery grant to RIC. Queen’s University is situated on traditional lands of the Anishinaabe and Haudenosaunee and the authors are grateful to conduct our research from these lands.

