## Supplementary Methods for "Evidence for continent-wide convergent evolution and stasis throughout 150 years of a biological invasion"

**This PDF includes:**

Supplementary Methods

Figs. S1 to S6

Tables S1 to S7

### Herbarium Samples

We used several methods to obtain GPS decimal coordinates, depending on the kind of location information provided on the specimen label and the database, as follows. We converted all available geographic coordinates into decimal WGS84 format. For herbarium specimens without GPS coordinates but with Public Land Survey System (PLSS) township and range codes given on the label, we converted the township-range-section codes into GPS coordinates through manually referencing Historic Map Works (http://www.historicmapworks.com), an online collection of historical documents. The online collection contains historical state atlases for counties and towns using PLSS codes as the coordinate system. We used Google Maps to manually georeference the latitudes and longitudes of township names found using the section number of the PLSS codes. For some specimens, GPS or PLSS coordinates could not be identified, but other landmark information was available, such as street names and intersections. For these records, we searched using Google Maps for location names hierarchically using the state before searching for the county within the state, and the township within the county. This was necessary because many county and township names are duplicated across US states. For specimens where the herbarium label gave the direction and distance of a collection site from a landmark, we used the measuring tool in Google Maps to determine the approximate location. Specimens that did not include sufficient information for geolocation were not included in the study.

### Weather Interpolation Scripts

*DownloadStationData.R* was written to scrape (i.e. download) local weather data for each collection site and calculate the length of the growing season to the time of collection in three basic steps. First, we identified up to 100 weather stations that (i) were active in the year of collection and (ii) were located within 0.5 degrees in latitude and longitude. Second, the stations were filtered if they were missing data, and only the closest twenty stations were included. Third, a vector of local growing temperature ($G_T$) was calculated for each weather station ($i$) for each day ($d$) in the collection year using the following formula:

$$G_{T_{i,d}} = \frac{T_{MAX_{i,d}} - T_{MIN_{i,d}}}{2} - 8$$

where $T_{MAX}$ is the highest temperature and $T_{MIN}$ is the lowest temperature recorded at a weather station $i$ on day $d$, except for days where $T_{MIN}$ is $<$ 8°C. The baseline temperature of 8°C was chosen based on previous studies (Shamsi & Whitehead 1974; Montague *et al.* 2008). Each $G_{T_i}$ is therefore a vector of length 365 (366 for leap years) with positive values indicating favorable growth conditions.

*CalculateSeasonMetricsIDW.R* uses the vectors $G_{T_i}$ from up to 20 weather stations closest to each herbarium specimen ($j$) to interpolate growing degree days by inverse distance weighting:

$$GDDs_{j,d} = \frac{\sum_i^n z_i \, G_{T_{i,d}}}{\sum_i^n z_i}$$

where $(n)$ is the number of stations, $GDDs_{i,d}$ is the interpolated growing-degree days for each day $(d)$ and weather station $(i)$, and $z_i = \frac{1}{x_i^2}$ where $x_i$ is the distance from the $i$th station to the collection site. Thus, $GDDs_{j,d}$ is a vector of 365 (or 366) cells containing values for daily mean temperatures interpolated from surrounding weather stations at each collection location. It is therefore possible to calculate $GDD_C$ and $GDD$ from the first day of the season to the day of collection, or the season end, respectively.

### Statistical Models

As described in the main text, we fit two different statistical models to ensure that the temporal trends were not an artifact of the chosen model.

### Table S1

**Table S1.** Number of herbarium specimens (total N = 3,429), separated by geographic region and sampling year.

| | A: <1900 | B: 1900-1939 | C: 1940-1980 | D: >1980 |
|---|---|---|---|---|
| East Coast | 95 | 445 | 822 | 418 |
| Midwest | 24 | 92 | 511 | 618 |
| West Coast | 1 | 18 | 122 | 263 |

### Table S2

**Table S2.** Mean estimates and 95% bootstrap confidence limits of the correlation between latitude and phenology index (z-score) for herbarium specimens and plants grown at one of three reciprocal transplant locations.

| Source | Mean | 95% Lower Bound | 95% Upper Bound |
|---|---|---|---|
| BEF | -0.866 | -0.949 | -0.748 |
| Herbarium | -0.702 | -0.922 | -0.167 |
| KSR | -0.771 | -0.901 | -0.600 |
| Timmins | -0.844 | -0.951 | -0.700 |

### Table S3

**Table S3.** Mean estimates and 95% bootstrap confidence limits of the correlation between season length and phenological development time ($\psi$), for herbarium specimens subdivided into three regions: East Coast, Midwest and West.

| Source | Mean | 95% Lower Bound | 95% Upper Bound |
|---|---:|---:|---:|
| East Coast | 0.6072 | 0.3396 | 0.8267 |
| Midwest | 0.7924 | 0.6169 | 0.8937 |
| West | 0.1692 | -0.1549 | 0.5003 |

### Table S4

**Table S4** Mean estimates and 95% bootstrap confidence limits of model coefficients from the nonlinear least squares model and logistic model predicting phenological index ($\varphi$).

| Term | Mean | 95% Lower Bound | 95% Upper Bound |
|---|---|---|---|
| *NLS model* | | | |
| $\beta_a$ | 2.04e-01 | 1.90e-01 | 2.19e-01 |
| $\beta_b$ | 6.49e-04 | 6.12e-04 | 6.89e-04 |
| $\beta_c$ | 9.07e-08 | 3.43e-08 | 1.52e-07 |
| $\beta_d$ | -1.18e-06 | -1.34e-06 | -1.02e-06 |
| $\beta_e$ | -3.16e-10 | -6.93e-10 | 1.69e-11 |
| *Logit model* | | | |
| (Intercept) | -0.81 | -1.06 | -5.33e-01 |
| SL | 1.09e-03 | -2.71e-03 | 4.16e-04 |
| SL:Age | 1.50e-06 | -2.10e-06 | 5.70e-06 |
| $GDD_c$ | 2.01e-03 | 1.72e-03 | 2.26e-03 |
| $GDD_c$:SL | 3.10e-06 | -4.40e-06 | -1.50e-06 |
| $GDD_c$:SL:Age | -5.10e-09 | -1.10e-08 | -7.63e-10 |
| $GDD_c$:Age | 1.40e-06 | 7.00e-07 | 2.30e-06 |
| Age | -6.13e-04 | -1.23e-03 | -8.93e-05 |

### Table S5

**Table S5.** Coefficients for a linear model estimating the effect of population age on the bootstrapped coefficient $\beta_{b,t}$ (Likelihood Ratio Test: $\chi^2$ = 8660.1, p < 0.001).

| | | Estimate | Std. Error |
|---|---|---|---|
| *Linear* | **(Intercept)** | 3.71e-04 | 2.47e-06 |
| | **age** | 3.13e-06 | 2.18e-08 |
| *Quadratic* | **(Intercept)** | 2.20e-04 | 2.80e-06 |
| | **age** | 7.30e-06 | 6.10e-08 |
| | **age^2** | -1.94e-08 | 2.73e-10 |

### Table S6

**Table S6.** Coefficients for a linear model estimating the effect of age on the bootstrapped coefficient $\beta_{d,t}$. (Likelihood Ratio Test: $\chi^2$ = 2154.8, p < 0.001)

| | | Estimate | Std. Error |
|---|---|---|---|
| *Linear* | **(Intercept)** | -3.442e-07 | 9.945e-09 |
| | **age** | -1.095e-08 | 8.772e-11 |
| *Quadratic* | **(Intercept)** | 1.495e-07 | 1.276e-08 |
| | **age** | -2.46e-08 | 2.778e-10 |
| | **age^2** | 6.339e-11 | 1.244e-12 |

### Figure S1

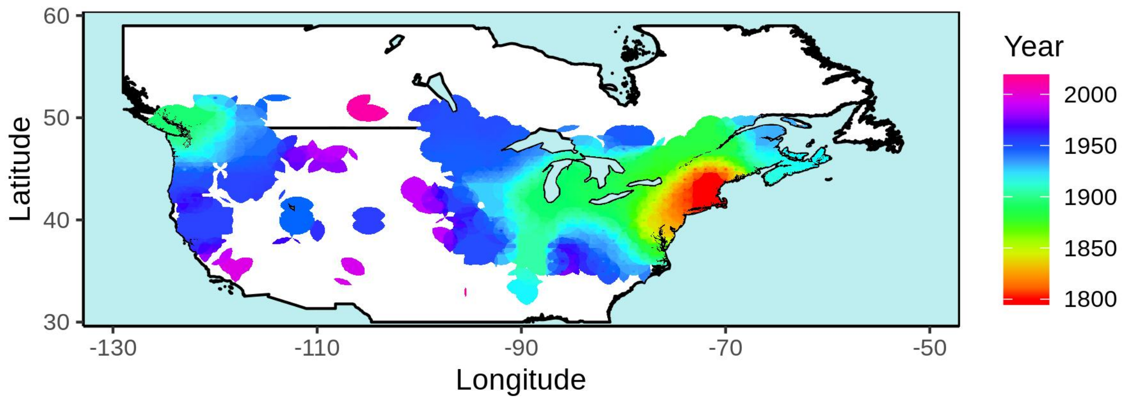


**Figure S1.** Estimated establishment dates across the North American distribution of *Lythrum salicaria*. Year of establishment was estimated by Kriging earliest recorded occurrences (from 11,735 records) across a grid of 1 degree latitude by 1 degree longitude.

### Figure S2

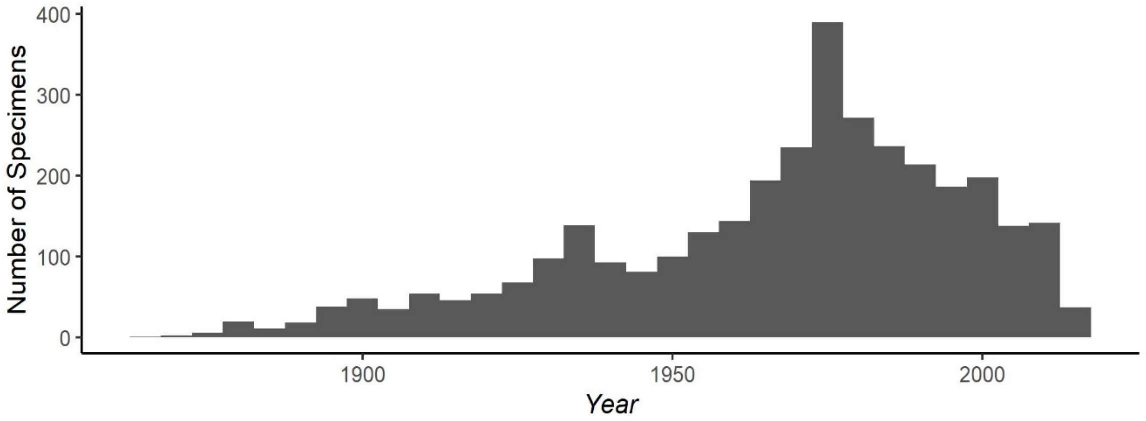


**Figure S2.** Histogram showing the number of *Lythrum salicaria* specimens collected by year from 1866 to 2016.

Figure S3

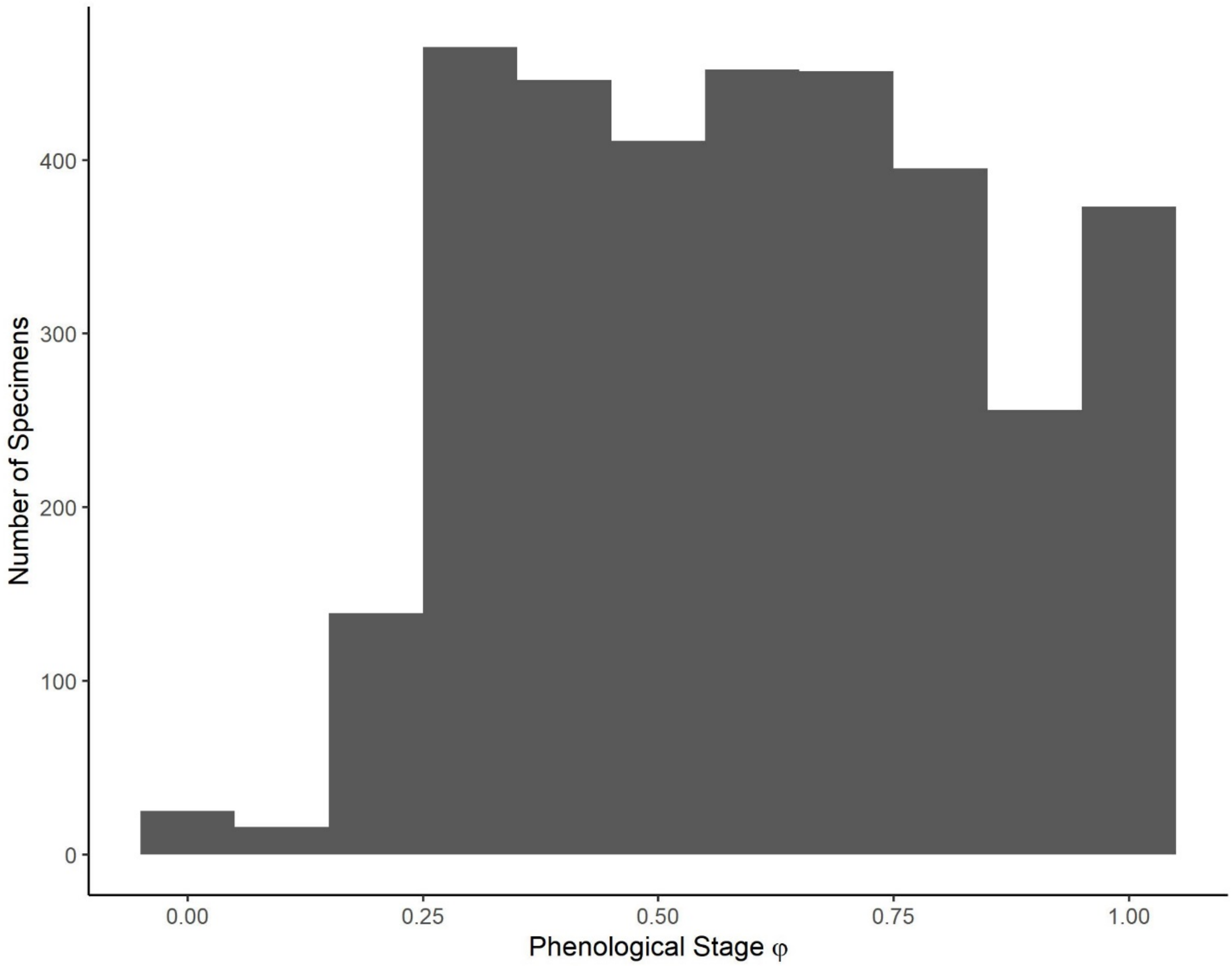


**Figure S3.** Distribution of phenological stage ($\varphi$) observed for each herbarium specimen, calculated from the relative lengths of buds, flowers, and fruits.

### Figure S4

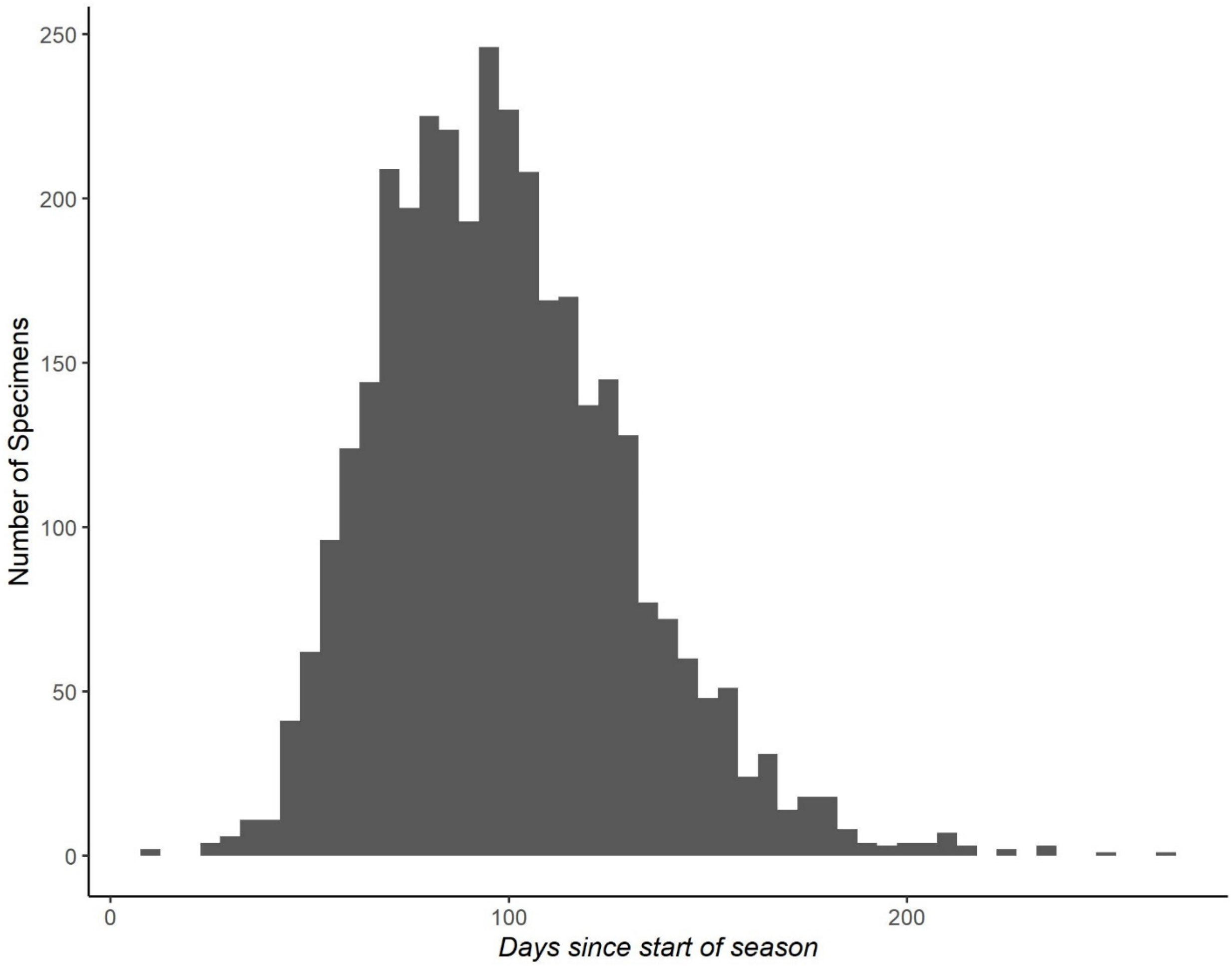


**Figure S4.** Histogram showing day of specimen collection relative to the calculated start of the growing season. Season start was identified as the first day after January 1 in an interval of 10 consecutive days in which mean temperature stayed above 8°C.

### Figure S5

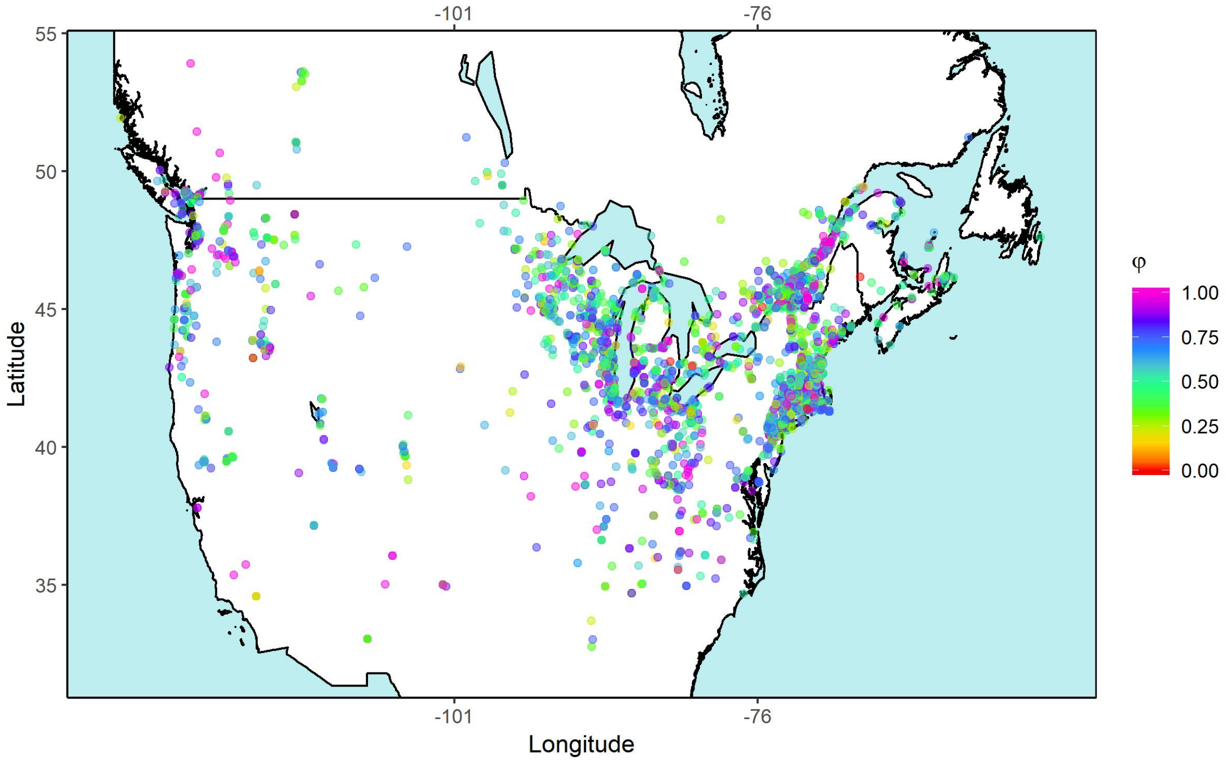


**Figure S5.** Map of 3,429 *Lythrum salicaria* herbarium specimens in North America, color-coded by phenological stage ($\varphi$).

Figure S6

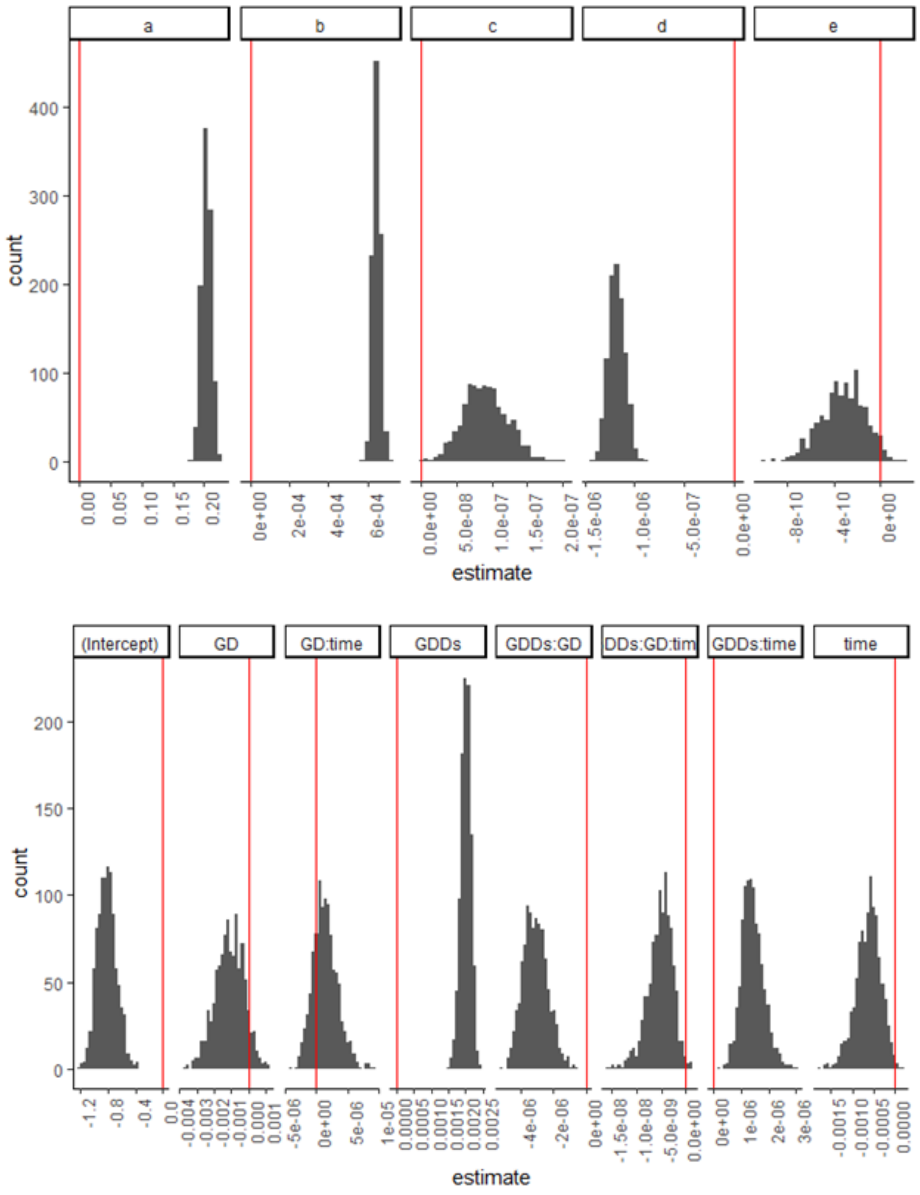


**Figure S6**. Bootstrapped parameter estimates of a Nonlinear Least Squares model (top row) and a logistical regression (bottom row) model. Model parameters for the NLS are provided in Eqs. 1 and 4 in the main paper. Model parameters for the logistic regression model are shown for the overall intercept, slope of season length (aka Growing Days or GD), growing degree days from the start of the growing season to the time of sampling (GDDs), time since population establishment (time) and interactions among the terms. Each bootstrap includes 1,000 spatially explicit resampling of herbarium records from a 0.1° latitude by 0.1° longitude grid.

### Figure S7

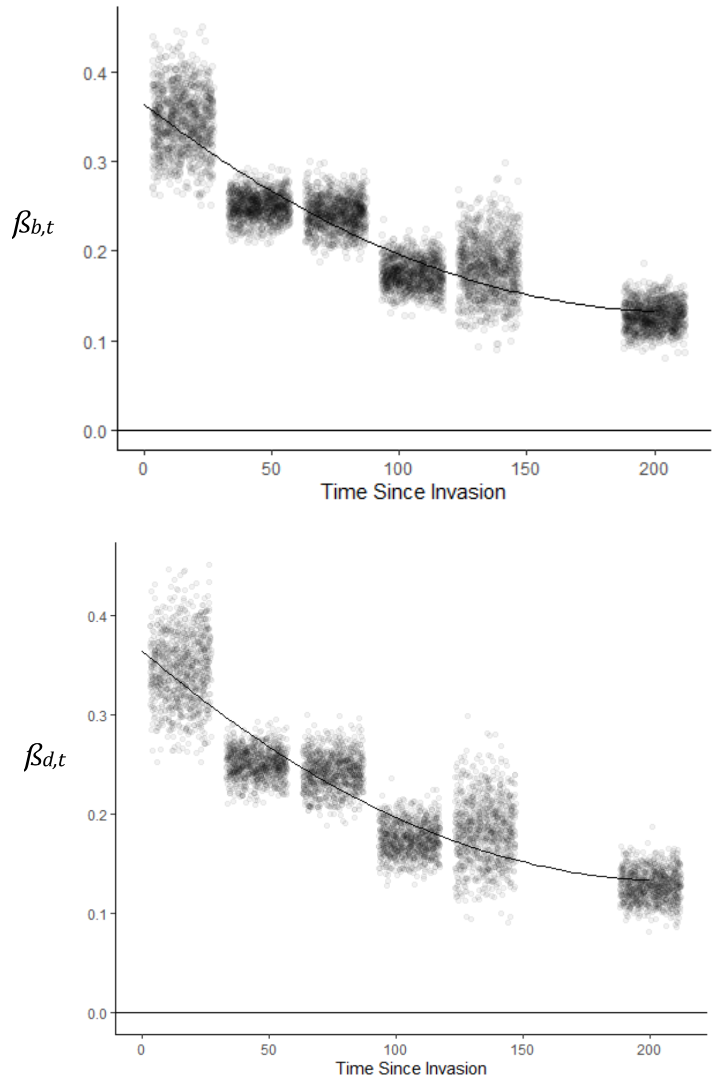


**Figure S7.** Bootstrap estimates and curve of best for of model coefficient from the Universal Logistic Model, separated for the Temperature Accumulation Coefficient ($\beta_{b,t}$, Top panel) and the Season Length Coefficient ($\beta_{d,t}$, Bottom Panel). Each panel includes the quadratic regression of best fit (LRT: $\chi^2 > 2158$, $P < 0.001$) and bootstrapped means from 1,000 spatially explicit resampling iterations (with replacement), binned into one of six classes of population age (i.e. time since invasion).
